# The atypical antidepressant and cognitive enhancer tianeptine disinhibits hippocampal CA1 pyramidal neurons by reducing GABAergic neurotransmission

**DOI:** 10.64898/2025.12.21.695760

**Authors:** SJ Martin, Y Li, B Tench, GD Phillips, JJ Lambert, D Belelli

**Affiliations:** Division of Neuroscience, School of Medicine, University of Dundee, MSI Building, Dundee, DD1 5HL, UK; Simons Initiative for the Developing Brain, University of Edinburgh, Hugh Robson Building, George Square, Edinburgh, EH8 9XF, UK

## Abstract

Tianeptine is an atypical antidepressant and cognitive enhancer whose actions are thought to be dependent on the activation of opioid receptors, including µ-opioid receptors expressed on hippocampal interneurons. Agonists of these receptors can cause a disinhibition of principal neurons by inhibiting GABA release or interneuron firing. However, direct evidence for a similar action of tianeptine has not previously been reported. We therefore studied the effects of tianeptine on in vivo and in vitro measures of inhibitory synaptic transmission. CA1 field excitatory postsynaptic potentials (fEPSPs) and local field potential (LFP) activity were recorded in area CA1 of anaesthetised rats. To obtain a more direct measure of tianeptine’s actions on GABA release, we also conducted whole-cell patch-clamp recordings of spontaneous and evoked inhibitory postsynaptic potentials (sIPSCs and eIPSCs) in CA1 pyramidal neurons in vitro. Intrahippocampal infusion of tianeptine (15 mM; 1 µl) caused a marked reduction in paired-pulse inhibition (PPI) of the population spike (20-ms inter-stimulus interval) elicited in the CA1 cell-body layer. Separate recordings in the stratum radiatum revealed an enhancement of beta and gamma-frequency LFP activity, an effect of tianeptine that we have reported previously. Infusion of the GABA_A_ receptor antagonist bicuculline caused a similar reduction in PPI but did not induce beta/gamma oscillations. In mouse hippocampal slices, tianeptine (50 µM) reduced both sIPSC frequency and eIPSC amplitude. These results confirm that tianeptine causes a disinhibition of CA1 pyramidal cells, but suggest that a simple blockade of GABAergic transmission is not alone sufficient to explain tianeptine’s effects on hippocampal network activity.

## Introduction

Tianeptine is an atypical antidepressant with memory enhancing and analgesic properties and an unusual mechanism of action. Despite its licensing for antidepressant use in France in 1989, and current approval in over sixty countries in Europe, Asia, and South America (Nishio et al., 2024), tianeptine’s mechanism of action remained uncertain until just over a decade ago. Although it has a tricyclic structure, tianeptine does not interact with noradrenaline or serotonin receptors, and does not inhibit the reuptake of these neurotransmitters (McEwen et al., 2010). In 2014, tianeptine was discovered to be an agonist of µ- and δ-opioid receptors (Gassaway et al., 2014), and subsequent work has revealed that µ-opioid receptor activation is necessary for its antidepressant effects (Samuels et al., 2017; Han et al, 2022), although an enhancement of AMPA-receptor-mediated transmission is also likely to be involved (Svenningsson et al., 2007; see Mariano et al., 2026). As a µ-OR agonist, tianeptine’s physiological effects at high doses resemble those of canonical opioids such as morphine, including undesirable side-effects such as respiratory depression (Hill et al., 2023), reduced gut motility (Sohn et al., 2012; Baird et al., 2022), and reward-related behaviours such as conditioned place preference (CPP; Han et al., 2022; Allain et al., 2023).

In the hippocampus, µ-ORs are prominently expressed in the somatodendritic regions and axon terminals of several subtypes of inhibitory GABAergic interneurons, particularly those targeting the somata of principal neurons (Svoboda et al., 1999; Drake & Milner, 1999 & 2002; Stumm et al., 2004; Won et al., 2023). Their activation causes a disinhibition of CA1 pyramidal cells (Zieglgänsberger et al., 1979; Neumaier et al., 1988; Lambert et al., 1991; McQuiston & Saggau, 2003; Shao et al., 2020), an action that is reminiscent of the opioid-induced disinhibition of dopaminergic neurons in the VTA (Bull et al., 2017). This disinhibition is likely mediated by the inhibition of voltage-gated calcium channels (VGCCs) leading to a reduction in GABA release (Capogna et al., 1993; Heinke et al., 2011), and / or the activation of potassium channels leading to the hyperpolarisation of interneurons and a reduction in their firing rate (Svoboda & Lupica, 1998; He et al., 2021); see Nam et al. (2021) for review.

We have previously reported that systemic administration of tianeptine causes beta-frequency oscillations in the hippocampus of anaesthetised rats (Burt et al., 2025), an action that might be related to the disinhibition of pyramidal neurons (cf. Gwilt et al., 2020). To address this possibility, we assessed the effects of tianeptine on in vivo and in vitro measures of the strength of GABAergic inhibition. First, we assessed the in vivo effect of local intrahippocampal tianeptine administration on the amplitude and paired-pulse inhibition (PPI) of the CA1 population spike, indirect measures of the strength of feedforward and feedback perisomatic inhibition that are reduced by µ-OR agonists in vitro (Kapur et al., 1989; Jedlicka et al., 2010; Giannopoulos & Papatheodoropoulos 2012). We then determined the effects of tianeptine on a more direct measure of synaptic inhibition using in-vitro whole-cell patch-clamp recordings of inhibitory postsynaptic currents (IPSCs). Specifically, we measured the frequency of spontaneous IPSCs (sIPSCs) and the amplitude of evoked IPSCs (eIPSCs), both of which are known to be reduced by the application of µ-OR agonists (e.g. Lupica, 1995; Shao et al., 2020). To determine whether the local hippocampal circuitry is sufficient for the generation of tianeptine-induced beta / gamma oscillations similar to those that we have previously observed after systemic administration, we examined the LFP changes resulting from intrahippocampal infusion of tianeptine in anaesthetised rats. Finally, we compared the in vivo actions of tianeptine on both PPI and LFP activity with those of locally applied bicuculline, a competitive antagonist of GABA_A_ receptors.

## Methods

### Animals

All procedures involving animals were conducted in accordance with the UK Animals (Scientific Procedures) Act (1986) and subject to local ethical review by the University of Dundee’s Welfare and Ethical Use of Animals in Research Committee. Female and male Lister hooded rats obtained from Charles River UK were used in all experiments. They were given at least 10 days to acclimatise before experiments were conducted. During this time, rats were housed in groups of 4 in pairs of cages, each 32 x 50 cm in area, and connected by an acrylic tube. They were given unrestricted access to food (standard raw chow, sunflower seeds, and wheat grains) and water, and maintained on a 12-h light / 12-h dark cycle at constant temperature (19-24°C). Bedding and nesting materials comprised wood shavings, paper wool, ‘sizzle-nest’, and hay, and enrichment items included acrylic tubes for refuge, wooden chew-sticks, and Aspen balls. For approximately 30 min per day, rats were placed, together with their cage-mates, in a ‘playpen’ measuring ∼0.75 x 1.5 m, and containing enrichment objects such as plastic tunnels, a running wheel, and a large cardboard refuge.

### Drugs

The vehicle for intrahippocampal infusion was artificial cerebrospinal fluid (aCSF). Drug solutions were dissolved in aCSF and frozen in small aliquots prior to use. The drugs and doses used were tianeptine hydrochloride (15 mM; Kemprotec Limited, Cumbria, UK) and (+)-bicuculline (10 µM, 30 µM, 0.1 mM, 0.5 mM, 1.5 mM, 15 mM; Sigma-Aldrich, Burlington, MA, USA. For in vitro experiments, a final concentration of 50μM tianeptine was prepared from a stock solution 500x higher in concentration.

### Surgery for in vivo fEPSP and LFP recording

At the start of each experiment, the rat was anaesthetised with 1.5 g/kg urethane (ethyl carbamate; 0.3 mg/ml; 5 ml/kg; IP) and placed in a stereotaxic frame (Kopf, Tujunga, USA) with the skull horizontal. Body temperature was monitored via a rectal probe and maintained at 36.2°C using a homeothermic heat blanket (Harvard Apparatus, Holliston, MA, USA). Depth of anaesthesia was assessed throughout the experiment, and 0.2 ml top-up doses of urethane were administered as required. Small holes were drilled in the skull for the insertion of the electrodes and cannulae. A small stainless steel jeweller’s screw was secured to the occipital bone to serve as a reference for fEPSP and LFP recording.

### In vivo recording of field excitatory postsynaptic potentials (fEPSPs)

To measure positive-going CA1 fEPSPs with superimposed population spikes, a recording electrode was placed in the stratum pyramidale (3.8 mm posterior and 2.5 mm lateral to bregma; depth approximately -2.1 mm from the dura) and a stimulation electrode was lowered into the contralateral stratum radiatum at a homotopic site (3.8 mm posterior and 2.5 mm lateral to bregma; depth approximately -2.4 mm from the dura) to recruit crossed Schaffer collateral / commissural projections (Fig. 1 A-C). Both stimulating and recording electrodes comprised paired twisted strands of PTFE-insulated platinum/iridium wire, external diameter = 0.103 mm. Correct electrode placement was verified by the characteristic depth profile of fEPSP changes observed during insertion; see Shires et al. (2012), Fig. 6, for a detailed explanation. Although both stimulating and recording electrodes were bipolar, only one of the two recording electrode channels was selected for analysis during each recording session.

**Fig. 1.**
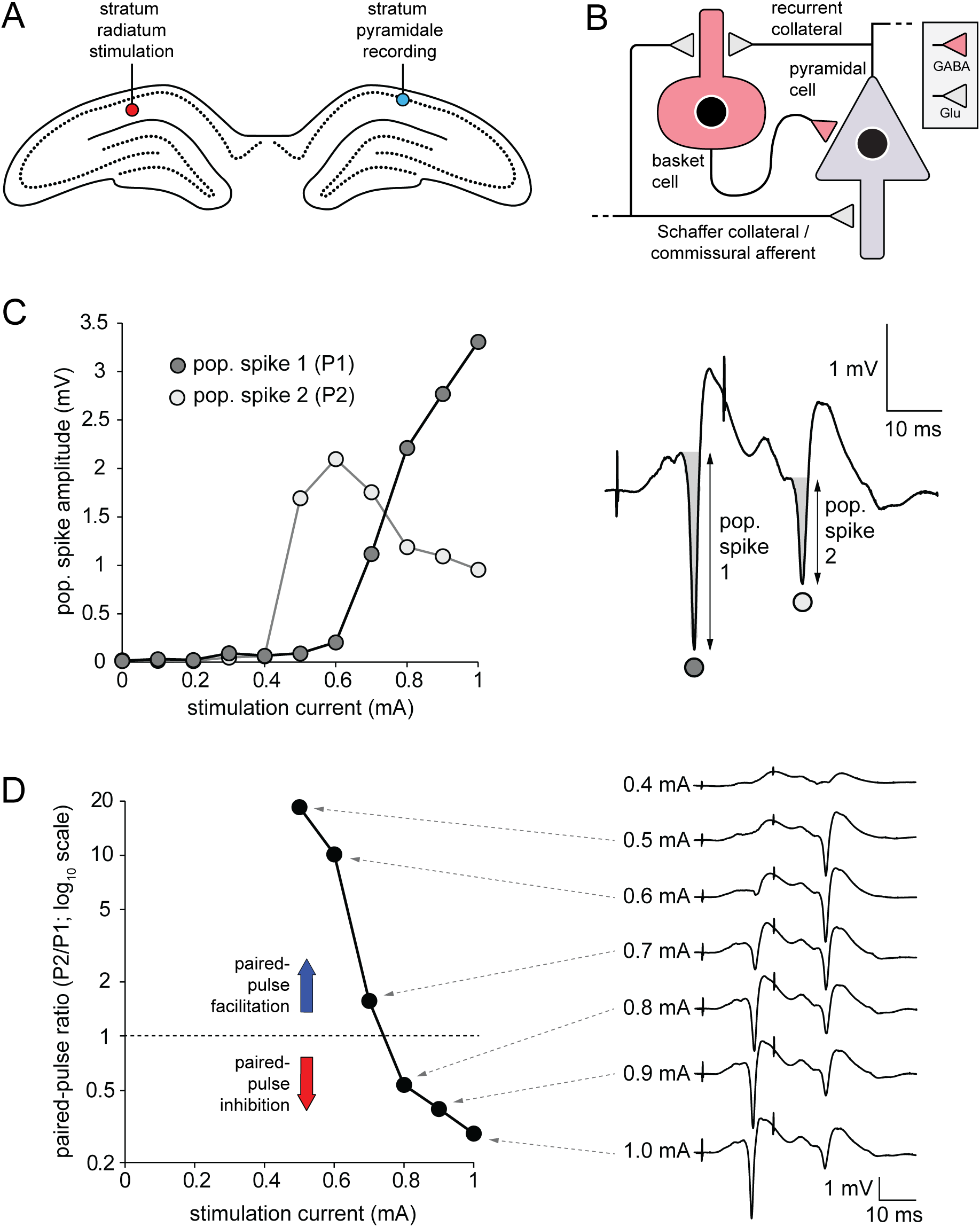
Protocol for inducing paired-pulse depression of the population spike (pop. spike) amplitude in area CA1 in vivo. (A) Location of recording electrode in the stratum pyramidale of area CA1 and stimulating electrode in the contralateral stratum radiatum. (B) Schematic figure of the stimulation of glutamatergic Schaffer collateral / commissural afferents and feedforward recruitment of an inhibitory basket cell. (C) Left-hand panel: dependence on stimulation intensity of the population spikes elicited by paired-pulse stimulation at a 50-ms inter-stimulus interval. Right-hand panel: example of the fEPSPs and population spikes elicited by paired stimulation at an intensity of 1.0 mA. Note the pronounced paired-pulse depression (PPD) of the second population spike. (D) Left-hand panel: Paired-pulse-depression ratio of the amplitude of the two population spikes as a function of stimulation intensity. Note the transition from paired-pulse facilitation to paired-pulse depression with increasing stimulation current.

Evoked fEPSPs were amplified and filtered (high pass = 0.3 Hz; low pass = 5 kHz) using a differential AC amplifier (model 3500, A-M Systems, Sequim, WA, USA) and sampled at 20 kHz using a data acquisition card (PCIe-6321; National Instruments, Austin, TX, USA) mounted in a PC running custom-written LabView software for the control of electrical stimulation and the time-locked recording of evoked fEPSPs (Evoked Potential Sampler, Patrick Spooner, University of Edinburgh). 50-Hz line noise was removed using a HumBug noise-elimination unit (Digitimer, Welwyn Garden City, UK). Stimulation was delivered via a Neurolog system and stimulus isolator units (Digitimer), and consisted of biphasic constant-current pulses, 100 µs per phase. At the start of each experiment, electrode depths were adjusted to maximise the amplitude of the positive-going fEPSP and negative-going population spike.

Cannulae for intrahippocampal infusion were constructed using 30-gauge insulin needles (external diameter = 0.31 mm) with bevelled tips, soldered to larger-gauge tubing and connected via plastic tubing to 5 µl SGE syringes mounted in a syringe pump (Model 11 Plus, Harvard Apparatus, Holliston, MA, USA). One syringe was always loaded with vehicle and one with tianeptine or bicuculline. Injection needles were placed bilaterally, 4.5 mm posterior and 3.0 mm lateral to bregma, with the needle tip -3.0 mm from the dura. In our experience, slow infusion rates and low volumes are required to avoid mechanical disturbance of the hippocampal tissue; for this reason, 1 µl of vehicle and drug solutions was administered over a 30-min period. To track the progress of infusion, a 0.5 µl air bubble was introduced into the infusion tubing. Movement of the bubble, and the absence of leakage at any point, was used to confirm the successful delivery of the vehicle or drug solution.

After electrode placement, paired stimulation pulses were delivered to measure PPI of the population spike amplitude. Stimulation intensity was adjusted until the amplitude of the second population spike was approximately 50% of the amplitude of the first (i.e. a paired-pulse ratio of 0.5). Intensities ranged from 0.5 – 1.0 mA, approximately double the intensity typically used for dendritic fEPSP recording (see Burt et al., 2025). Baseline recording comprised paired-pulse stimulation at 20-s intervals; once stable fEPSPs and population spikes had been recorded for at least 40 min, a vehicle infusion (1 µl of aCSF over 30 min) was delivered via the cannula ipsilateral to the recording electrode. After the end of infusion, recording continued for a further 30 min. At the end of this period, the former recording electrode was lowered into the stratum radiatum, both poles were connected to the stimulus isolator units to allow stimulation, and the former stimulating electrode was raised into the stratum pyramidale, and its outputs were connected to the differential amplifier. This allowed the recording of positive-going fEPSPs and population spikes from the hippocampus contralateral to the one previously used for vehicle infusion. As before, stimulation intensity was adjusted until the amplitude of the second population spike was approximately 50% of the amplitude of the first, and baseline recording began. Once stable fEPSPs and population spikes had been recorded for at least 40 min, an infusion of tianeptine (1 µl of15 mM over 30 min) or bicuculline (1 µl of 10 µM, 30 µM, 0.1 mM, 0.5 mM, 1.5 mM, or 15 mM over 30 min) was delivered, followed by a further 30-min recording period. The hemisphere receiving drug versus vehicle infusion (i.e. left versus right) was counterbalanced across treatment groups and sex according to a randomised block design.

For each EPSP, the amplitude of the population spike was calculated by measuring the voltage difference between the negative peak of the population spike and the preceding local maximum (Fig. 1C; right-hand panel). The paired-pulse ratio was obtained by dividing the amplitude of the second population spike by the amplitude of the first.

Values were then re-normalised to the mean paired-pulse ratio over the 10 min prior to the start of infusion, so that the baseline paired-pulse ‘ratio’ was assigned a value of 1 for each recording. The first and second population spikes were individually normalised to their mean amplitudes over the 10 min before the start of infusion. This baseline was assigned a value of 100%, and subsequent changes were expressed as a percentage of the baseline value. For each measure, mean values were calculated over successive 2-min recording periods for the duration of the experiment.

### In vivo spontaneous LFP recording

For LFP recording, stimulating and recording electrodes were positioned exactly as described above, except that both electrodes were placed in the stratum radiatum of CA1 (c. -2.4 mm from the dura bilaterally). Correct electrode placement was verified by the characteristic depth profile of fEPSP changes observed during insertion, and final electrode depths were adjusted to maximise the amplitude of the negative-going fEPSP elicited by contralateral stimulation. Although single stimulation pulses (biphasic constant-current pulses, 100 µs per phase) were delivered at 20-s intervals throughout the experiment to elicit fEPSPs, subsequent analysis focussed primarily on the continuous wide-band LFP that was recorded in parallel via the same recording electrode. Although the recording electrode was bipolar, only one of the two recording electrode channels was selected for LFP recording and analysis during each recording session.

Spontaneous LFP activity was amplified and filtered (high pass = 0.3 Hz; low pass = 5 kHz) using a differential AC amplifier (see above) and sampled using a USB-6003 acquisition device (National Instruments, Austin, TX, USA) connected to a PC running custom-written software for LFP capture (Patrick Spooner, University of Edinburgh). Data were acquired at 20 kHz, low-pass filtered at 100 Hz and down-sampled to 200 Hz. Successive 2-s samples of data (50% window overlap; temporally filtered using the Hann function) were analysed by fast Fourier transform (FFT). Samples containing fEPSPs were omitted from the analysis. Mean power spectral density from 1-80 Hz was analysed in 0.5 Hz frequency bins and averaged across successive 2-min recording periods. Data were converted to log_10_ values. Time-frequency plots were constructed by plotting successive 2-min power spectra as a 3-D heat map with power as the z-axis. The change in spectral power before and after drug injection was calculated by subtracting the log_10_ power at each time point and frequency bin from the corresponding mean over the 10-min period before injection, then plotted as a 3-D time-frequency graph, this time with change in spectral power relative to baseline as the z-axis. The mean change in power was also calculated in specific frequency bands including beta (10-30 Hz) and slow gamma (30-50 Hz).

### Histology

In a subset of those experiments in which cannulae were implanted (n = 8), brains were removed at the end of the experiment and post-fixed in 10% formalin. Subsequent histological processing, sectioning, and scanning was carried out by the Dundee Imaging Facility at the University of Dundee. The fixative was first removed, and brains were washed 3 times with 1X phosphate-buffered saline. Brains were then placed in 30% sucrose in PBS (pH = 7.4) overnight at 4 °C, or until the tissue sank, then removed and embedded in OCT, and frozen on dry ice. The region of the dorsal hippocampus containing the infusion cannulae was sectioned at 30 µm, at -20°C, using a Leica CM1860UV cryostat (Leica Microsystems (UK) Ltd., Sheffield, UK). Slide-mounted sections were air dried for 1-2 hours prior to storage at -20 °C and staining with haematoxylin and eosin (H&E). Slides were scanned at 20X using a Zeiss Axioscan 7 (Carl Zeiss Ltd., Cambourne, UK).

### In vitro IPSC recording

Adult male C57Bl/6J mice (3-4 weeks old) were killed by a Schedule 1 method, in accordance with UK Home Office and institutional guidelines. The brain was rapidly dissected and incubated in an ice-cold, oxygenated, artificial cerebrospinal fluid (aCSF) sucrose-based solution, which contained (in mM): sucrose (234), glucose (10), NaHCO_3_ (26), NaHPO_4_ (1.25) MgSO_4_ (10), KCl (2.5) CaCl_2_ (0.5). The solution was bubbled with 95% O_2_ / 5% CO_2_ (pH = 7.4). Sagittal 350-μm-thick hippocampal brain slices were cut using a Leica VT1200 vibratome (Leica Microsystems (UK) Ltd., Sheffield, UK). Following slicing the tissue was maintained at room temperature in an extracellular solution of the following composition (in mM): NaCl (126), NaHCO_3_ (26), KCl (2.9), NaH_2_PO_4_ (1.25) MgCl_2_ (2), CaCl_2_ (2), glucose (10), and bubbled with 95% O_2_ / 5% CO_2_ (pH = 7.3).

After at least 1 h under these conditions, whole-cell voltage-clamp recordings of GABAergic transmission were made from CA1 pyramidal neurons. An Olympus BX51 (Olympus, Southall, UK) microscope, equipped with differential interference contrast infrared (DIC/IR) optics, was used to visually identify CA1 pyramidal neurons in the pyramidal-cell layer, located in close apposition to the border of stratum radiatum. The brain slices were continuously perfused at room temperature (26-27 °C) with the extracellular solution described above, bubbled with 95% O_2_ / 5% CO_2_ (pH = 7.3), but now additionally supplemented with kynurenic acid (2mM) to inhibit ionotropic glutamate receptors. Voltage-clamp recordings (Vh = -40 to -60) of evoked and spontaneous IPSCs (eIPSCs, sIPSCs respectively) were performed using an Axopatch 1D amplifier (Molecular Devices, Union City, CA, USA). The recording pipette (3-4 MΩ) was fabricated from borosilicate glass tubing with a 2.0-mm outer diameter and 0.5-mm wall thickness (Garner Glass Co., Claremont, CA, USA) using a P83 puller (Narashige, Tokyo, Japan). The internal pipette solution contained (in mM): CsCl (135), HEPES (10), EGTA (10), CaCl_2_ (1), MgCl_2_ (1), ATP (2), QX314 (5) (pH = 7.3). For electrical stimulation of GABAergic synaptic inputs, a 4–6 MΩ stimulating pipette filled with extracellular solution was placed in the stratum radiatum layer very close to the border with the stratum pyramidale and within 50-100 μM of the recorded pyramidal neurons, using a stimulation interval of 0.1 s, with a stimulation intensity of 20-60 μA and duration of 60 μs.

All recordings were analysed offline using the Strathclyde Electrophysiology Software (Electrophysiology Data Recorder (WinEDR) and Whole Cell analysis Program (WinWCP); courtesy of Dr J. Dempster, University of Strathclyde). Individual sIPSCs were detected using a low amplitude (–4 pA, 3 ms duration) threshold detection algorithm followed by visual scrutiny to avoid spurious detections. Analysis of the sIPSCs was restricted to events with a rise time ≤ 1 ms to minimize the contribution of dendritically generated currents, which are subject to cable filtering. Individual accepted events were analysed for peak amplitude and 10–90% rise time. For each cell the peak amplitude and rise time of a minimum of 100 (range 106-332) sIPSCs was determined for both the control and the drug condition (after 10 min of tianeptine perfusion). sIPSC frequency analysis was performed by sampling and averaging 3 x 10-s segments across a 2-min period both during the baseline and 10 min after tianeptine application. For the eIPSCs resulting from electrical stimulation of the stratum radiatum layer, the fast rise time inclusion criterion was not employed (although the RT was > 1ms in only one cell that had an RT of 1.5 ms), and the mean peak amplitude was determined for a minimum of 22 events (range 22-58) for both the baseline period and in the presence of tianeptine.

Numerical data were analysed using Microsoft Excel and IBM SPSS (IBM, Armonk, New York, USA). Graphs were prepared using Excel, SigmaPlot (Grafiti LLC, Palo Alto, CA, USA), and Adobe Illustrator (Adobe Systems, San Jose, CA, USA). Data are displayed as the arithmetic mean ± standard error of the mean (SEM), and individual data points are shown when bar graphs are presented. Comparisons were conducted using 2-tailed Student’s paired-sample t-tests, and correlational analyses were conducted by calculating Pearson’s correlation coefficient (r) and the corresponding p-value.

## Results

In anaesthetised rats, paired-pulse stimulation of the contralateral Schaffer collateral / commissural projection (Fig. 1A & B) yielded a biphasic effect on the amplitude of the second population spike recorded in the CA1 stratum pyramidale, depending on the stimulation intensity applied. Examples from a single rat are shown in Fig. 1C & D. Paired-pulse facilitation was observed at moderate stimulation intensities (e.g. 0.5 mA), but a pronounced paired-pulse inhibition (PPI) occurred at higher intensities (e.g. 1.0 mA). In subsequent experiments, we chose a stimulation intensity that yielded a paired-pulse ratio of ∼0.5, i.e. a 50% reduction in amplitude between first and second population spikes.

We first examined the effects of local intrahippocampal infusion of tianeptine on population spike amplitude and PPI in CA1 stratum pyramidale (Fig. 2A). In each animal, vehicle infusion was carried out in one hippocampus, before the electrodes were re-positioned for recording in the contralateral hippocampus during tianeptine infusion (see Materials and Methods for full details). Infusion of tianeptine caused a gradual increase in the amplitude of the first and second population spikes elicited by stimulation of Schaffer collateral / commissural afferents in the contralateral stratum radiatum (Fig. 2 B-D). An increase in paired-pulse ratio was also evident, reflecting a proportionally greater effect on the amplitude of the second population spike relative to the first (Fig. 2E). These effects were typically maximal at, or soon after, the end of the 30-min infusion period. Infusion of vehicle caused no pronounced changes in these measures (open circles in Fig. 2C-E). Analysis of the mean values 20-30 min after the start of infusion revealed a significantly larger increase in the first population spike following tianeptine versus vehicle administration [Fig. 2C; right-hand panel; t(3) = 3.59; p = 0.037; paired-samples t-test], but a non-significant (though numerically larger) difference in the second population spike [Fig. 2D; right-hand panel; t(3) = 2.30; p = 0.11]. The paired-pulse ratio was also significantly higher after tianeptine versus vehicle treatment [Fig. 2E; right-hand panel; t (3) = 3.30; p = 0.046; paired-sample t-test].

**Fig. 2.**
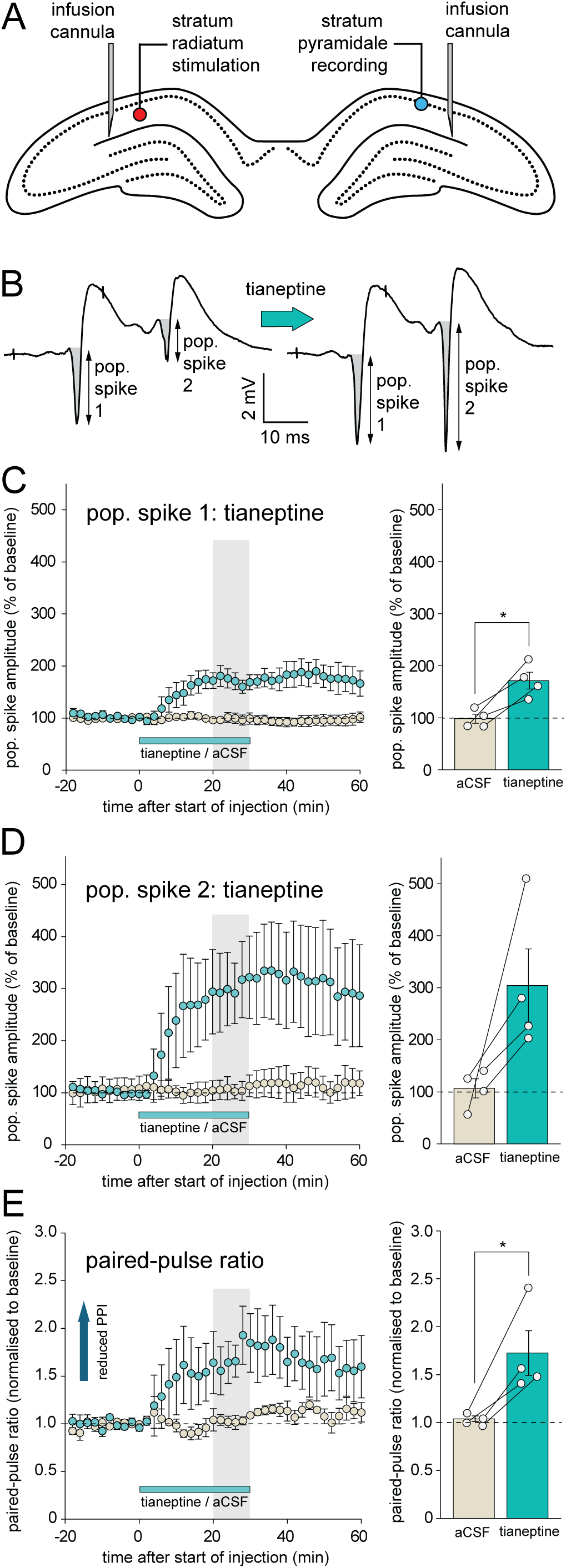
Tianeptine (1 µl; 15 mM) causes a reduction in paired-pulse depression in vivo (n = 4). (A) Bilateral placement of injection cannulae adjacent to recording and stimulating electrodes in the stratum pyramidale and contralateral stratum radiatum. (B) Examples of fEPSPs and population spikes elicited by paired-pulse stimulation before (left-hand side) and after (right-hand-side) the infusion of tianeptine adjacent to recording site. Note the slight increase in the amplitude of population spike 1, and the marked reduction in the paired-pulse inhibition of population spike 2. (C) Left-hand panel: time course of the increase in the amplitude of population spike 1 during and after vehicle or tianeptine infusion. Right-hand-panel: mean population spike 1 amplitude 20-30 min after the start of vehicle or tianeptine infusion (grey bar in left-hand panel) (*p < 0.05; paired-sample t-test). (D) Increase in the amplitude of population spike 2 during and after infusion; other details as for C. (E) Left-hand panel: time course of the change in paired-pulse ratio after infusion; right-hand panel: mean paired-pulse ratio 20-30 min after the start of infusion (*p < 0.05; paired-sample t-test).

To obtain a more direct measure of tianeptine’s impact on GABAergic neurotransmission, we conducted whole-cell patch clamp recordings from CA1 pyramidal neurons in the stratum pyramidale of mouse hippocampal slices. A stimulation electrode was placed close to the patch electrode to recruit perisomatic interneurons (Fig. 3A). There was no change in the mean amplitude of sIPSCs 10 min after application of tianeptine relative to baseline values [Fig. 3B; t(5) = 0.65; p = 0.54; n= 6 slices from 3 mice; examples of individual sIPSCs are shown in the upper panel]. However, the mean frequency of sIPSCs recorded over a 2-min period before and 10 min after application of tianeptine was significantly reduced [Fig. 3C; t(5) = 7.06; p = 0.0009; n = 6 slices from 3 mice; representative examples of a 1-s recording period before and after tianeptine application are shown in the top panel. Fig. 3D shows the effects of tianeptine on eIPSCs elicited by electrical stimulation close to the patch electrode. Tianeptine caused a significant decrease in peak eIPSC amplitude [Fig. 3D, left-hand panel [t(5) = 3.22; p = 0.024], and peak amplitude values expressed as a percentage of baseline were significantly below 100% after tianeptine application [Fig. 3D, right-hand panel; t(5) = 5.47; p = 0.0028; one-sample t-test; 6 slices from 3 mice; representative examples of individual eIPSCs before and after tianeptine application are shown in the upper panel].

**Fig. 3.**
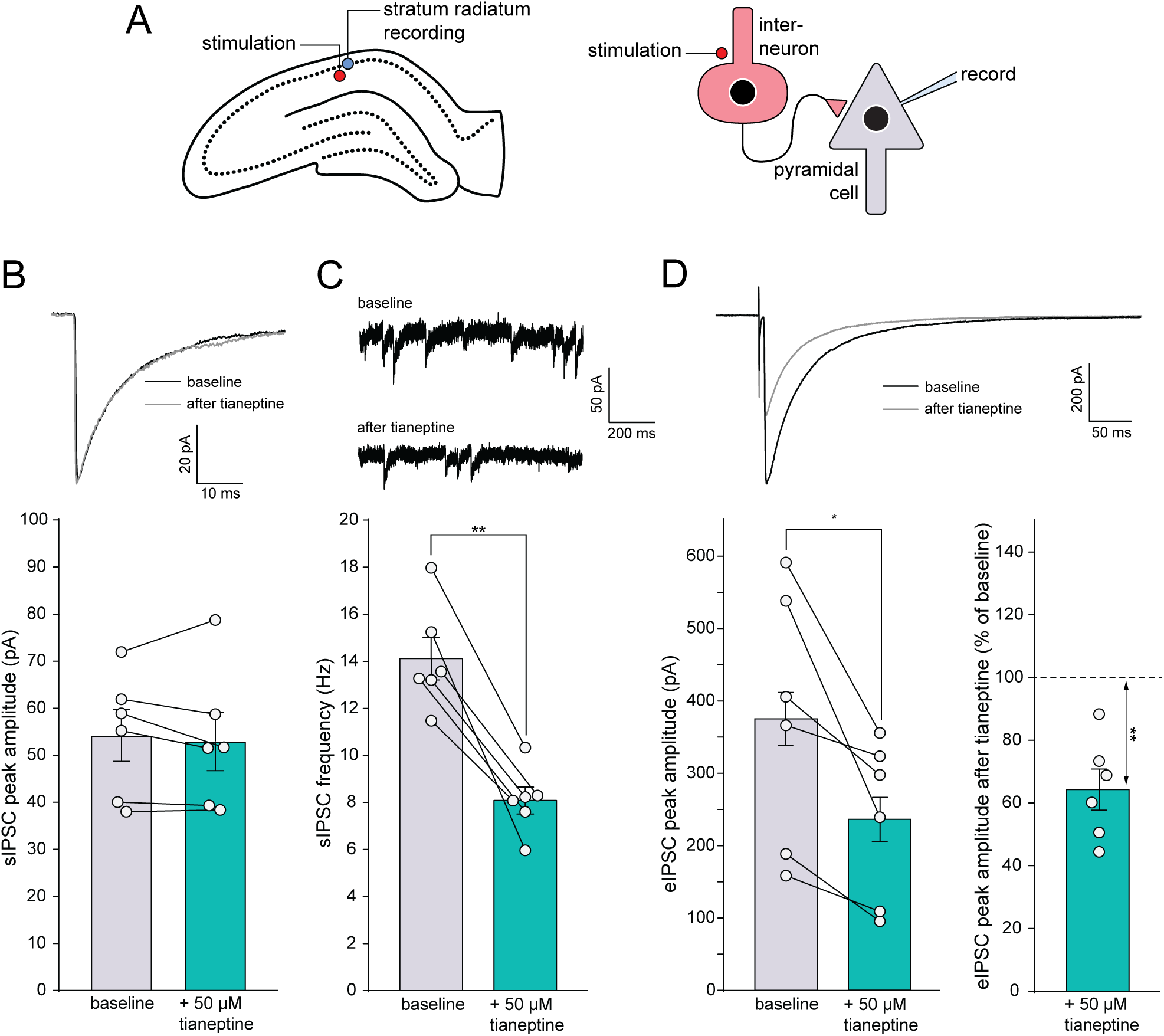
Tianeptine (50 µM) reduces spontaneous and evoked IPSCs in CA1 pyramidal neurons in vitro. (A) Location of patch electrode in the CA1 stratum pyramidale and an adjacent stimulating electrode to recruit inhibitory interneurons. (B) Upper panel: examples of sIPSCs recorded before and after the application of tianeptine; lower panel: mean change in sIPSC peak amplitude (n = 6 slices from 3 mice). (C) Upper panel: one-second segments of recordings from a single neuron illustrating a reduction in the frequency of sIPSCs after tianeptine application; lower panel: mean change in sIPSC frequency (*p < 0.05; paired-sample t-test; n= 6 slices from 3 mice). (D) Upper panel: examples of eIPSC amplitude before and after tianeptine application; lower panel: mean change in eIPSC amplitude after tianeptine application (n = 6 slices from 3 mice), expressed as raw values (left; *p < 0.05; paired sample t-test) and percentage of baseline (right; **p < 0.01; one-sample t-test).

To determine the effects of local intrahippocampal infusion of tianeptine on hippocampal network activity, we carried out additional in vivo experiments in which spontaneous LFP activity was recorded from the stratum radiatum as in Burt et al. (2025). The bilateral placement of infusion cannulae and recording electrodes is shown in Fig. 4A (see Materials and Methods for full details). Infusion of tianeptine caused a pronounced increase in beta and gamma-frequency LFP activity, whereas little change in power was evident after vehicle infusion (Fig. 4C-H). Analysis of the time-course of beta (10-30 Hz; Fig. 4I) and slow gamma (30-50 Hz; Fig. 4J) activity showed a peak increase in power during the last 10 min of infusion. Analysis of the mean change in power 20-30 min after the start of infusion revealed a significant difference between tianeptine and vehicle administration in both beta [Fig. 4I; right-hand panel; t(3) = 4.16; p = 0.025; paired-sample t-test] and slow gamma ranges [Fig. 4J; right-hand panel; t(3) = 6.47; p = 0.0075].

**Fig. 4.**
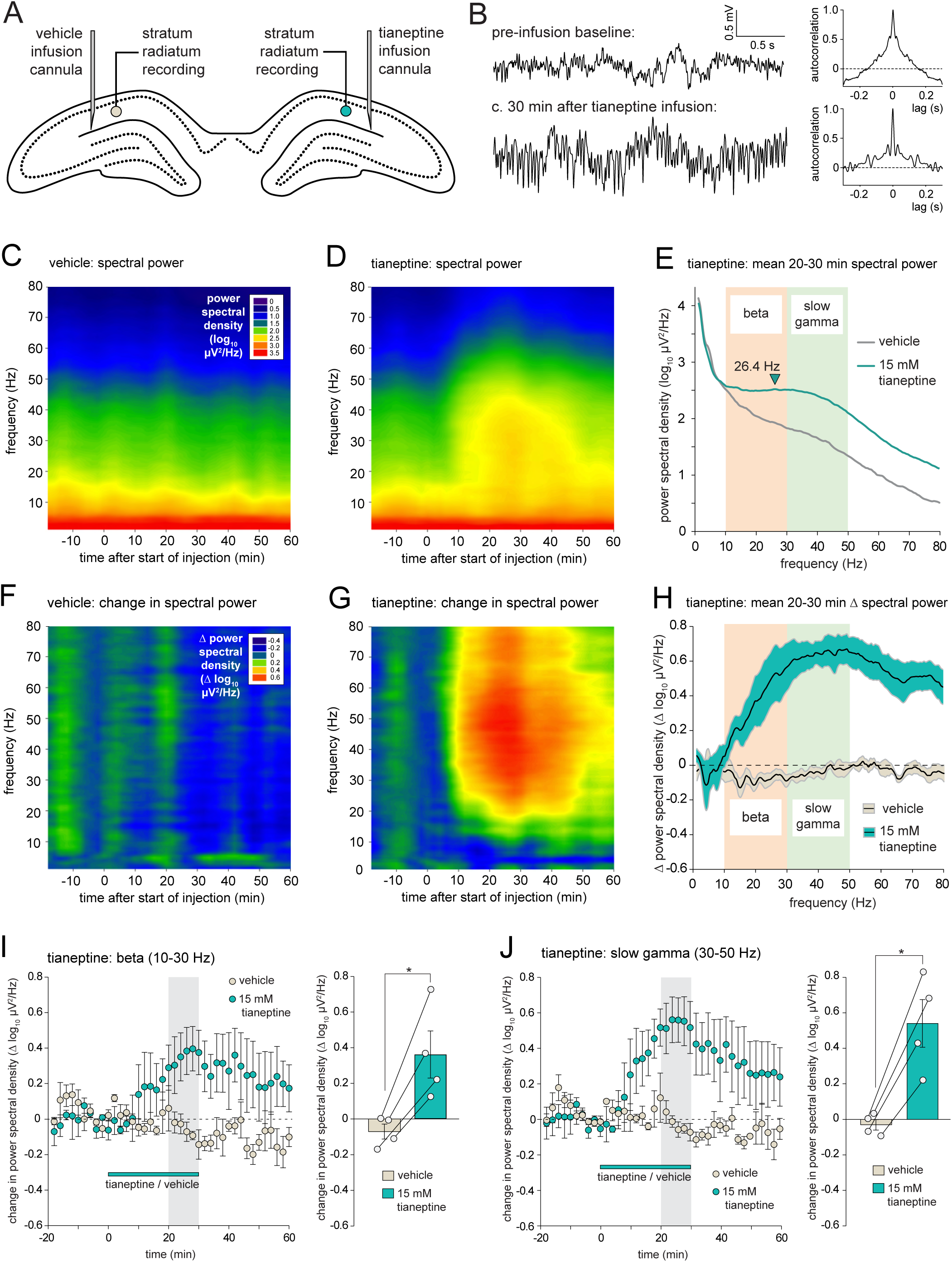
Tianeptine (1 µl; 15 mM) causes an increase in hippocampal beta / gamma oscillations in vivo (n = 4). (A) Bilateral placement of infusion cannulae and recording electrodes in the stratum radiatum. In reality, the hemispheres containing drug and vehicle cannulae were counterbalanced across animals. (B) Samples of LFP activity recorded from a single rat before and ∼30 min after the start of tianeptine infusion (i.e. at the end of the infusion period), indicating a pronounced increase in rhythmic beta / low gamma. (C) Time-frequency plot of spectral power after vehicle infusion. (D) Time-frequency plot of spectral power after tianeptine infusion; z-scale as in C. (E) Mean spectral power 20-30 min after the start of infusion in vehicle and tianeptine hemispheres; error bars omitted for clarity. (F) Change in spectral power after vehicle infusion. (G) Change in spectral power after tianeptine infusion; z-scale as in F. (H) Mean change in spectral power 20-30 min after the start of vehicle and tianeptine infusion. The black lines indicate the means, and the filled areas bounded by grey lines indicate ± 1 SEM. (I) Left-hand panel: time-course of changes in beta-frequency power normalised to the pre-infusion baseline; right-hand panel: mean change in beta-frequency power 20-30 min after the start of vehicle and tianeptine infusion (*p < 0.05; paired-sample t-test). (J) Change in slow-gamma-frequency power; other details as for I.

In the next experiment, we compared the effects of tianeptine on in vivo LFP and PPI measures with the effects of directly blocking GABA_A_ receptors with the competitive antagonist bicuculline. To obtain a dose of bicuculline that reduced inhibitory transmission without causing epileptiform activity, we first conducted a dose-response study of the effects of six different concentrations of bicuculline on CA1 stratum radiatum fEPSPs and LFP activity. Fig. 5A shows the effects of increasing doses of bicuculline on fEPSP slope; bars represent data from individual animals, except for the vehicle bar that indicates the mean of six vehicle infusions that carried out prior to bicuculline administration in the contralateral hemisphere in each animal. Marked decreases in fEPSP slope were evident only at the highest dose administered, 1 µl of 15 mM bicuculline infused over 30 min. Fig. 5B shows individual examples of fEPSPs before and ∼30 min after the start of bicuculline infusion. There was little change in the fEPSP after 30 µM bicuculline infusion, aside from the appearance of a small positive-going population spike. However, at a dose of 15 mM bicuculline, fEPSPs were markedly prolonged and multiple population spikes were evident.

**Fig. 5.**
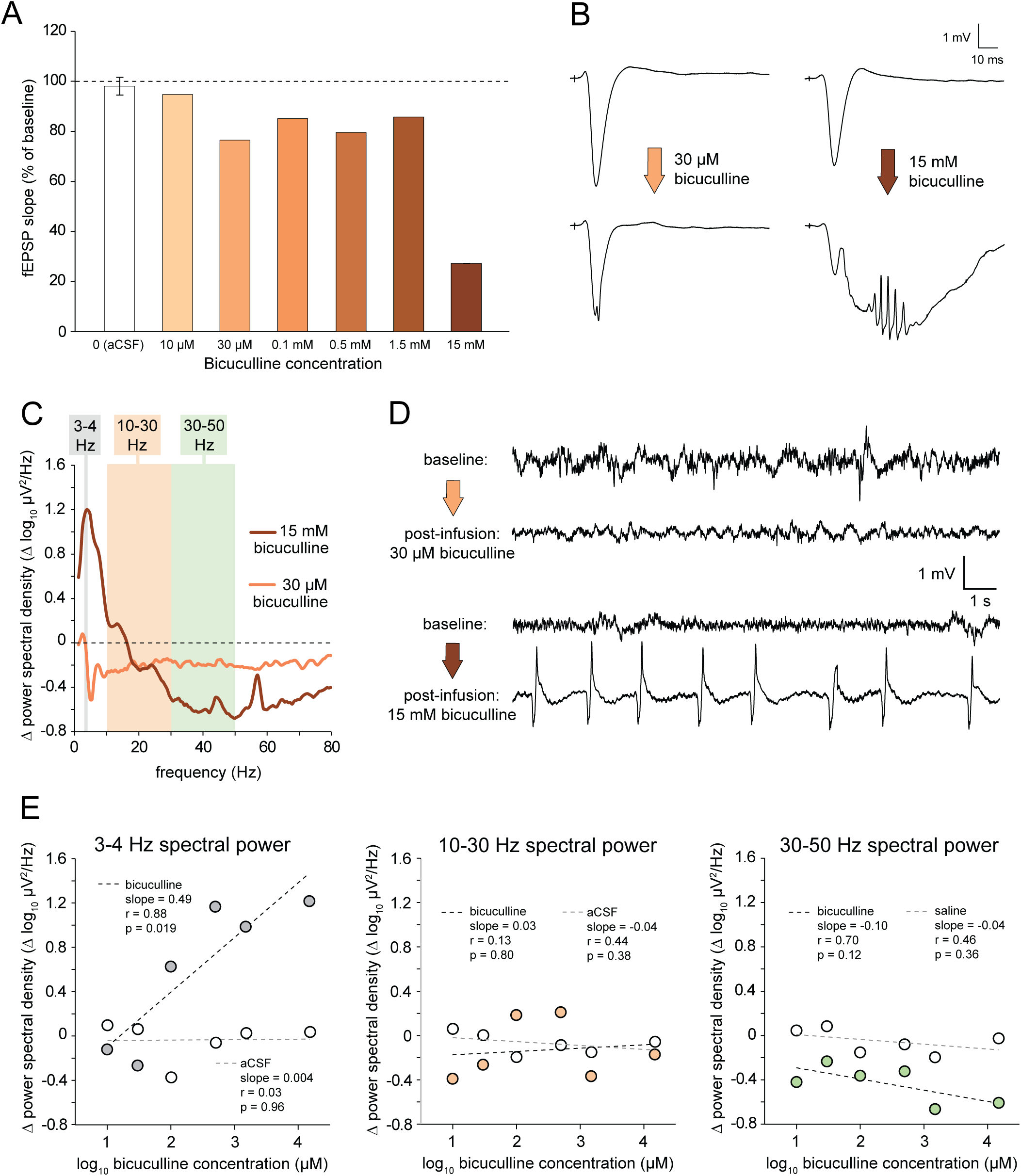
Dose-dependent effects of GABA_A_ receptor antagonism on CA1 evoked fEPSPs and spontaneous LFP activity in CA1 in vivo. (A) Reduction in fEPSP slope caused by infusion of increasing doses of bicuculline. (B) Examples of fEPSPs recorded before and after the infusion of 30 µM and 15 mM bicuculline. Note the appearance of epileptiform activity comprising multiple population spikes at the higher dose. (C) Power spectral changes 20-30 min after the start of infusion of 30 µM and 15 mM bicuculline. (D) Examples of raw LFP recordings before and c. 30 min after the start of infusion of 30 µM and 15 mM bicuculline. (E) Correlation between mean spectral power in each of three frequency bands 20-30 min after infusion and bicuculline concentration (filled circles); data from the corresponding vehicle-infused hemisphere are shown in each case (open circles).

We next analysed continuous LFP activity from the same recordings. Fig. 5C shows mean spectral power 20-30 min after infusion of 30 µM and 15 mM bicuculline. The latter dose caused an increase in low-frequency LFP power with a peak at ∼4 Hz, but a broad decrease in beta and gamma power. In contrast, 30 µM bicuculline caused a small drop in spectral power that did not change markedly as a function of frequency. Fig. 5D (upper traces) shows samples of LFP activity recorded before and ∼30 min after infusion of 30 µM bicuculline, illustrating the decrease in amplitude of both low and high-frequency components of activity. The lower traces show samples of LFP activity before and after 15 mM bicuculline. After infusion, LFP activity was dominated by large amplitude spontaneous LFP spikes reminiscent of interictal spikes. Although the frequency composition of these events caused a mean spectral peak in the theta-frequency range of 3-4 Hz, the pattern of LFP activity produced by 15 mM bicuculline was distinct to the rhythmic 3-4 Hz theta activity sometimes observed under urethane anaesthesia (see Martin et al., 2019; Fig. 5). Fig. 5E shows the mean change in 3-4 Hz spectral power 20-30 min after bicuculline infusion as a function of dose (grey circles). There was a significant correlation between the change in spectral power and the log concentration of bicuculline (r = 0.89; n = 6; p = 0.019). An increase in 2-4 Hz power was observed at all concentrations above 30 µM. For example, infusion of 0.1 mM bicuculline (log_10_ concentration in µM = 2 on the x-axis), there was an increase in spectral power of 0.63 log units, corresponding to a 4.25-fold increase in power. This was associated with the appearance of large amplitude spontaneous LFP events similar to those illustrated in Fig. 5D. This pattern of activity grew more pronounced with increasing doses of bicuculline. The open circles in Fig. 5E (left-hand panel) show the change in spectral power after vehicle infusion (prior to the infusion of bicuculline) in each of the six animals tested. Vehicle infusion had little effect on spectral power, and, as expected, there was no significant correlation between the change in spectral power and the dose of bicuculline that was later infused into the contralateral hippocampus of each animal (r = 0.03; n = 6; p = 0.38). There was no significant relationship between bicuculline concentration and the change in beta-frequency (10-30 Hz) power (Fig. 5E; middle panel) or slow-gamma power (Fig. 5E; right-hand panel). However, in the latter frequency band, a paired-sample t-test comparing vehicle and bicuculline data at each concentration revealed an overall dose-independent fall in slow gamma power induced by bicuculline [t(6) = 6.40; p = 0.0014; paired-samples t-test].

Since 30 µM bicuculline was the highest concentration that did not result in large amplitude spontaneous LFP events, we selected this dose for further investigation and comparison with the effects of tianeptine. We first studied the in vivo effects of local intrahippocampal infusion of this dose on population spike amplitude and PPI recorded in CA1 stratum pyramidale (Fig. 6A). As before, vehicle infusion was first carried out in one hippocampus, before the electrodes were re-positioned for recording in the contralateral hippocampus during bicuculline infusion (see Materials and Methods for full details). Like tianeptine, bicuculline caused an increase in the amplitude of the first population spike (Fig. 6A), and numerically larger increase in the second population spike (Fig. 6B), and in increase in paired-pulse ratio (Fig. 6E). Analysis of the mean values 20-30 min after the start of infusion revealed a significantly larger increase in the first population spike following bicuculline versus vehicle administration [Fig. 6C; right-hand panel; t(3) = 4.79; p = 0.017], and a trend towards a difference in the second population spike [Fig. 6D; right-hand panel; t(3) = 2.97; p = 0.0592; paired-sample t-test]. The paired-pulse ratio was also significantly higher after bicuculline treatment [Fig. 6E; right-hand panel; T93) = 3.99; p = 0.043].

**Fig. 6.**
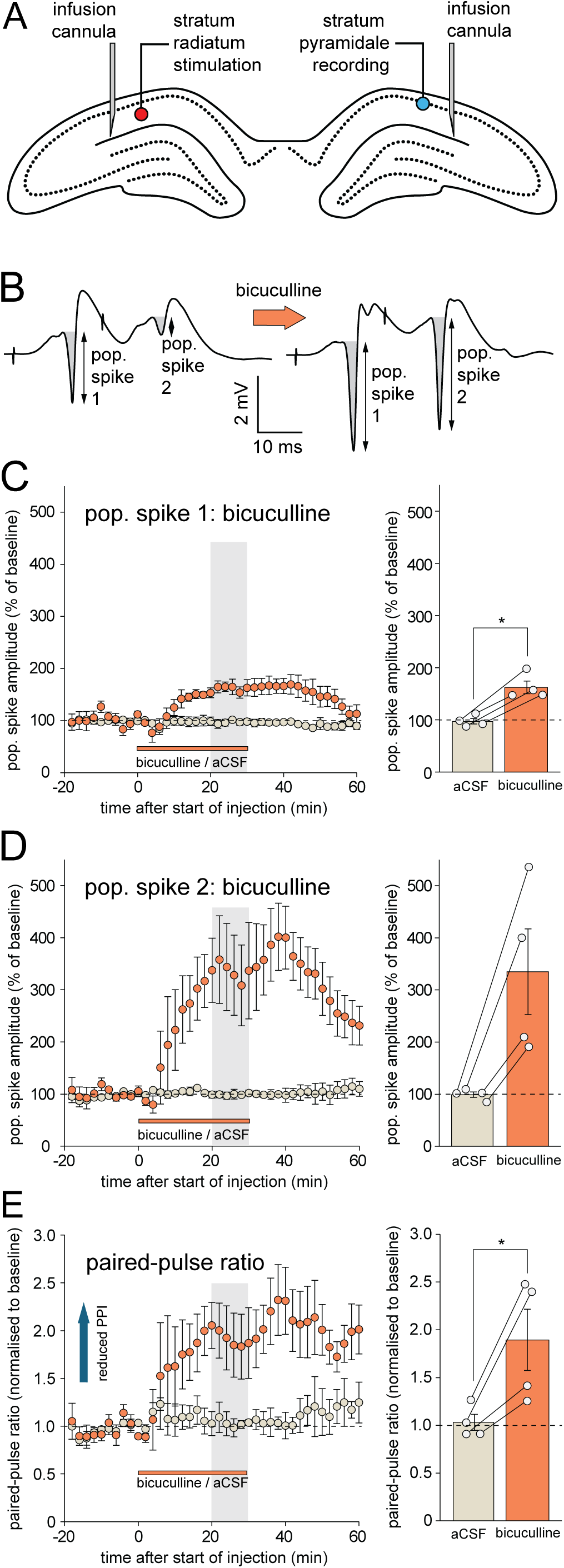
Bicuculline (1 µl; 30 µM) causes a reduction in paired-pulse depression in vivo (n = 4). (A) Bilateral placement of injection cannulae adjacent to recording and stimulating electrodes in the stratum pyramidale and contralateral stratum radiatum. (B) Examples of fEPSPs and population spikes elicited by paired-pulse stimulation before (left-hand side) and after (right-hand-side) the infusion of bicuculline adjacent to recording site. Note the increase in the amplitude of population spike 1, and the marked reduction in the paired-pulse inhibition of population spike 2. (C) Left-hand panel: time course of the increase in the amplitude of population spike 1 during and after vehicle or bicuculline infusion. Right-hand-panel: mean population spike 1 amplitude 20-30 min after the start of vehicle or bicuculline infusion (grey bar in left-hand panel) (*p < 0.05; paired-sample t-test). (D) Increase in the amplitude of population spike 2 during and after infusion; other details as for C. (E) Left-hand panel: time course of the change in paired-pulse ratio after infusion; right-hand panel: mean paired-pulse ratio 20-30 min after the start of infusion (*p < 0.05; paired-sample t-test).

As with tianeptine, we carried out parallel experiments in which spontaneous LFP activity was recorded from the stratum radiatum during bicuculline infusion. The bilateral placement of infusion cannulae and recording electrodes is shown in Fig. 7A (see Materials and Methods for full details). Infusion of bicuculline caused a slight drop in LFP power across the entire frequency range (Fig. 7B-H), consistent with the dose-response data shown in Fig. 5. Analysis of the change in power in beta (10-30 Hz) and slow gamma (30-50 Hz) bands revealed significantly lower beta power 20-30 min after the start of bicuculline versus vehicle infusion (Fig. 7I; t(5) = 3.20; p = 0.024; paired-sample t-test). This difference did not reach significance in the slow-gamma band (Fig. 7J; t(5) = 2.12; p = 0.088; paired-samples t-test).

**Fig. 7.**
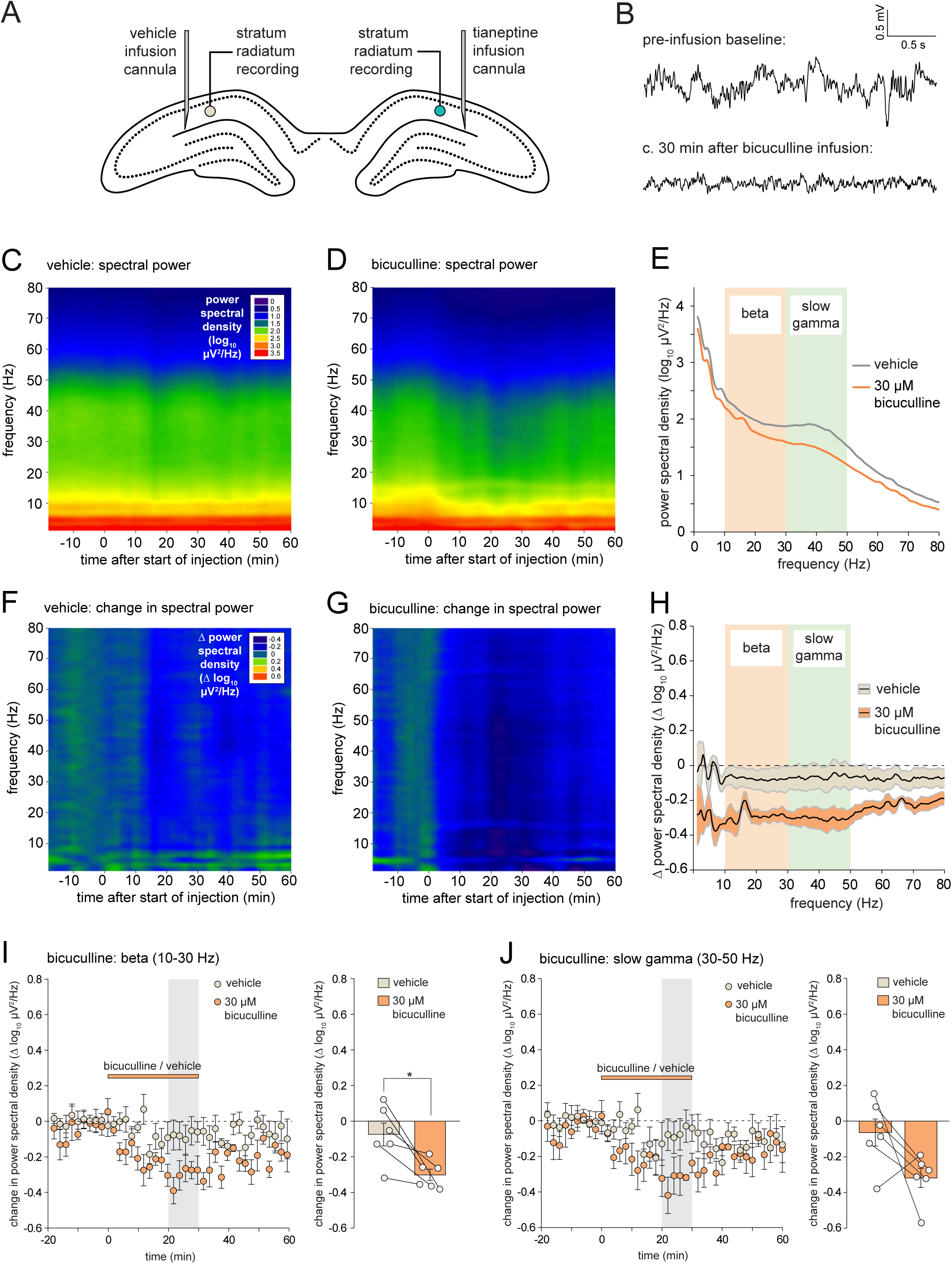
Bicuculline (1 µl; 30 µM) causes a modest broad-spectrum decrease in hippocampal LFP power in vivo (n = 4). (A) Bilateral placement of infusion cannulae and recording electrodes in the stratum radiatum. (B) Samples of LFP activity recorded from a single rat before and c. 30 min after the start of bicuculline infusion, indicating a modest decrease in the amplitude of LFP activity. (C) Time-frequency plot of spectral power after vehicle infusion. (D) Time-frequency plot of spectral power after bicuculline induction; z-scale as in C. (E) Mean spectral power 20-30 min after the start of infusion in vehicle and bicuculline hemispheres; error bars omitted for clarity. (F) Change in spectral power after vehicle infusion. (G) Change in spectral power after tianeptine infusion; z-scale as in F. (H) Mean change in spectral power 20-30 min after the start of vehicle and tianeptine infusion. The black lines indicate the means, and the filled areas bounded by grey lines indicate ± 1 SEM. (I) Left-hand panel: time-course of changes in beta-frequency power normalised to the pre-infusion baseline; right-hand panel: mean change in beta-frequency power 20-30 min after the start of vehicle and tianeptine infusion (*p < 0.05; paired-sample t-test). (J) Change in slow-gamma-frequency power; other details as for I.

Fig. 8A shows an example of an H&E-stained section showing the location of an infusion cannula in the left hippocampus. In all brains examined, infusion cannulae were correctly situated, typically in area CA1 close to the hippocampal fissure (Fig. 8B).

**Fig. 8.**
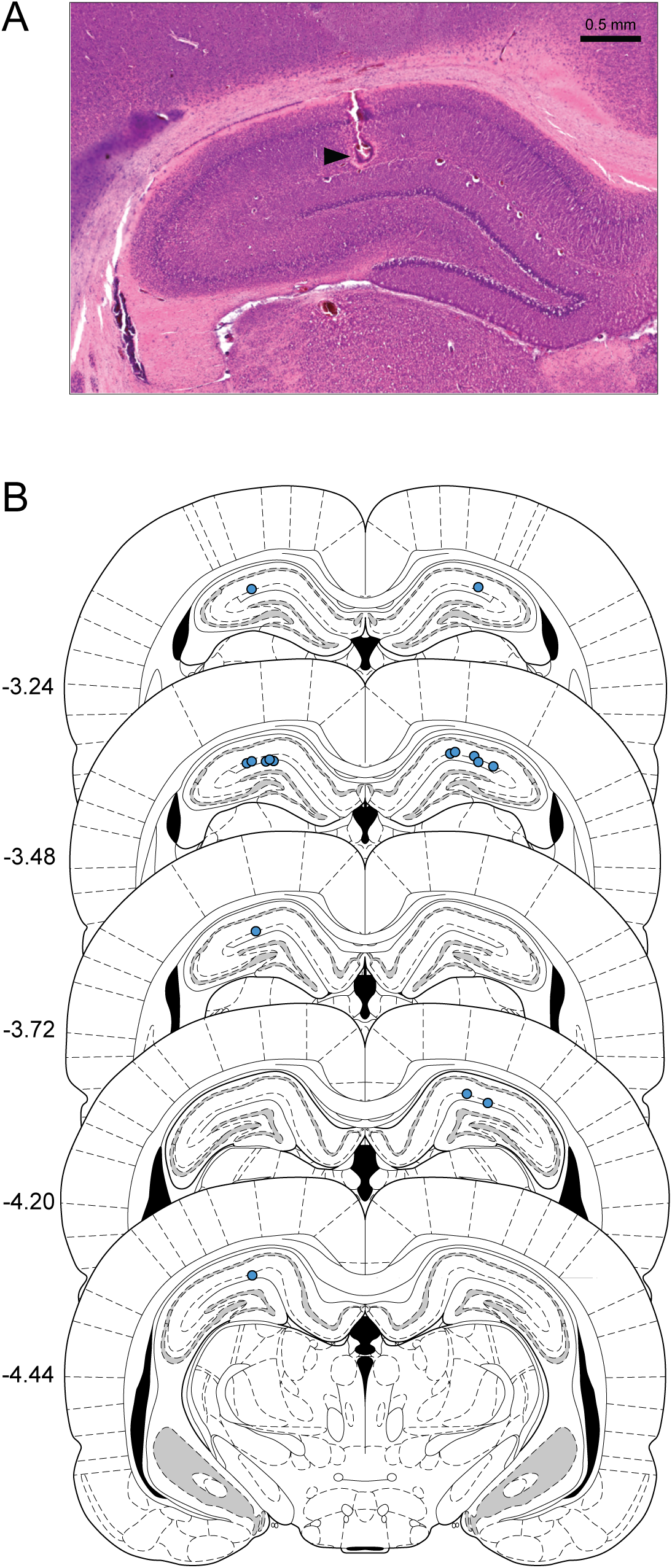
Infusion sites. (A) Example of an H&E-stained section showing the infusion site at the tip of the cannula track. (B) Infusion locations in the subset of brains that were sectioned and stained. Cannula positions are marked on the corresponding coronal section from the Paxinos and Watson (2005) atlas. All cannulae were correctly sited in the hippocampus.

## Discussion

The central finding of this study is that tianeptine, an atypical antidepressant and cognitive enhancer, causes a disinhibition of dorsal hippocampal CA1 pyramidal neurons. In urethane-anaesthetised rats, local intrahippocampal infusion of tianeptine caused a reduction in paired-pulse inhibition of the CA1 population spike, an indirect measure of the strength of synaptic inhibition. In whole-cell recordings of CA1 pyramidal cells in vitro, tianeptine caused a reduction in the frequency but not amplitude of spontaneous IPSCs, and a reduction in the amplitude of IPSCs evoked by electrical stimulation, providing more direct evidence for a reduction in the release of GABA from inhibitory interneurons. We also observed that local intrahippocampal administration of tianeptine in vivo causes hippocampal beta/gamma-frequency oscillations, following on from our previous report of an increase in beta-frequency LFP power after systemic injection (Burt et al., 2025). Local blockade of GABA_A_ receptors via the intrahippocampal infusion of bicuculline caused a similar reduction in PPI to tianeptine but did not increase LFP power in the beta- or gamma-frequency ranges.

Since the discovery that tianeptine is an opioid receptor agonist, many of its physiological and behavioural effects have been attributed to its actions on this receptor class (e.g. Han et al., 2022), and we have previously reported that tianeptine causes hippocampal beta-frequency LFP oscillations that are blocked by the opioid receptor antagonist naloxone (Burt et al., 2025). Our preliminary data from awake mice indicate that similar oscillations are observed in wild-type animals but absent in µ-OR knockout mice (Trigo et al., 2022). The µ-OR receptor is also likely to be responsible for the increase in excitability and reduction in PPI of the CA1 population spike observed in the current study. Increases in CA1 excitability and population spike amplitude have been reported in many studies; for an early review of this phenomenon, see Siggins et al. (1986). Indeed, application of morphine or DAMGO causes a reduction in CA1 PPI in brain slices (Linseman & Corrigall, 1982; Giannopoulos & Papatheodoropoulos, 2013).

While µ-ORs are found on several interneuron subtypes in the hippocampus, high levels of expression are found in almost all parvalbumin-positive (PV+) basket cells (Svoboda et al., 1999; Drake & Milner, 1999 & 2002; Won et al., 2023), fast-spiking neurons that make perisomatic synaptic contacts with CA3 and CA1 pyramidal cells, and closely regulate their output (Hijazi et al., 2023). The recurrent or feedforward activation of these cells by the collaterals of principal neuron axons is thought to play a central role in PPI of the population spike (Sloviter et al., 1991; Jedlicka et al., 2010). An inhibition of GABA release mediated by the activation of mu opioid receptors expressed on these neurons likely underlies tianeptine’s ability to increase excitability and reduce PPI.

However, tianeptine is also an agonist of δ-ORs, though with lower potency relative to µ-ORs (Gassaway et al., 2014). δ-ORs are expressed on several types of hippocampal interneurons, including PV+ cells (Erbs, 2012; He et al., 2021), but application of an agonist selective for δ-ORs does not reduce CA1 paired-pulse inhibition in response to Schaffer collateral stimulation in vitro (Lupica & Dunwiddie, 1991), instead decreasing feedforward inhibition mediated by the activation of temporoammonic projections to CA1 (Rezai et al., 2013). The δ-OR is therefore unlikely to be responsible for tianeptine’s suppression of PPI in the current experiment.

A more direct index of disinhibition is provided by whole-cell patch-clamp recording of IPSCs in CA1 pyramidal neurons, a measure of phasic, synaptic inhibition. Our observation of a reduction in the frequency of spontaneous IPSCs and the amplitude of electrically evoked IPSCs after tianeptine administration suggests a reduction in the vesicular release of GABA, and is consistent with previous reports of similar effects after the application of agonists selective for µ-ORs (Lambert et al., 1991; Lupica et al., 1992; Lupica, 1995; McQuiston, 2008; Shao et al., 2020). A likely mechanism is the activation of µ-ORs located on axon terminals—these suppress presynaptic neurotransmitter release by inhibiting voltage-gated calcium channels (Weiss & Zamponi, 2021). However, activation of µ-ORs located on the soma or dendrites of the same neurons can reduce neuronal excitability and hence firing rate by increasing the activation of a variety of potassium channels (Svoboda & Lupica, 1998). Either of these mechanisms could underlie the IPSC changes observed in the present study, as well as the enhancement of PPI in vivo. However, the observation that miniature IPSCs recorded in the presence of tetrodotoxin are reduced in frequency after administration of DAMGO (Lupica, 1995) indicates that a direct suppression of GABA release is sufficient to cause disinhibition, even if other actions of the µ-OR play a role under more physiological conditions.

As with suppression of PPI observed in vivo, PV+ interneurons are likely mediators of tianeptine’s reduction of GABA release. Evidence for this is provided by a recent study in which DAMGO caused a profound reduction in the amplitude of IPSCs elicited by selective optogenetic activation of CA1 PV+ interneurons; however, a more modest but still significant reduction was observed after conventional electrical stimulation that presumably recruited a mixed population of interneuron types (Shao et al., 2020). In the current study, the stimulating electrode was placed close to the recording electrode, ∼100 µm superficial to the pyramidal cell layer in the proximal stratum radiatum. This is likely to be an effective location for the recruitment of perisomatic inhibition by neurons including PV+ basket cells.

We have previously observed beta-frequency oscillations after systemic administration of tianeptine (Burt et al., 2025), but in this past study all brain areas expressing µ- and possibly δ-ORs will have been affected by the drug. The observation that systemic drug administration causes LFP oscillations in a specific structure does not mean that the oscillations are locally generated—they may be dependent on phasic or patterned activity originating elsewhere and conveyed by long-range afferents. There is good evidence that tianeptine has physiological actions at many different sites in the brain. For example, tianeptine’s locomotor stimulant effects are dependent on the activation of µ-ORs on NAc medium spiny neurons (MSNs) expressing dopamine D1 receptors, and its antidepressant effects require the activation of µ-ORs on ventral hippocampal SST+ interneurons (Han et al., 2022). Although this has not been investigated for tianeptine itself, other µ-OR agonists cause a disinhibition of dopamine neurons and increased release of dopamine in the nucleus accumbens (NAc) (e.g. Bull et al., 2017). VTA dopamine neurons, as well as glutamatergic and GABAergic afferents (Kim et al., 2024) also innervate the hippocampus—although the majority of dopaminergic afferents target the ventral part of the structure (Takeuchi, Duszkiewicz et al., 2016; Tse, Privitera et al., 2023), rather than the dorsal hippocampus studied in the current experiment. Other potential upstream targets include the entorhinal cortex (Gurgenidze et al., 2022), medial septum / diagonal band of Broca (Alreja et al., 2000), and the nucleus reuniens (Jayachandran et al., 2023; de Mooij-van Malsen, 2023). Nonetheless, the observation of beta- and gamma-frequency oscillations occurring very soon after local administration of tianeptine in the present study suggests that the hippocampal circuitry alone is sufficient for the generation of these rhythms, even if other brain areas play additional or modulatory roles after systemic injection.

Naturally occurring hippocampal gamma oscillations originate from multiple sources, including CA3 and the entorhinal cortex, with ‘slow gamma’ (∼30-50 Hz) typically originating in CA3 (Colgin, 2015). Although tianeptine increased spectral power over a wide range of beta and gamma frequencies in the current study, the largest change lay within the slow-gamma range, consistent with a site of action in CA3. The broader frequency composition of tianeptine-induced oscillations in the current study might reflect the absence of tianeptine induced effects in upstream regions that drive or modulate hippocampal activity. However, we also cannot rule out the possibility that the current results reflect higher tianeptine concentrations at the recording electrode after local versus systemic administration.

The mechanism by which tianeptine enhances gamma oscillations remains a puzzle, although similar effects have been reported with other µ-OR receptor agonists such as buprenorphine (Burt et al., 2025) and morphine (Reakkamnuan et al., 2015; Sakae & Martin, 2019). In area CA3, gamma oscillations can arise from mutually connected populations of excitatory pyramidal cells and inhibitory interneurons, particularly fast-spiking parvalbumin-positive (PV+) interneurons such as basket cells that mediate perisomatic inhibition and control the timing of pyramidal cell firing (for review, see Mann et al., 2005; Bartos et al., 2007; Buzsaki & Wang, 2012; Colgin, 2016), though other sub-types such as SST+ interneurons have also been implicated (e.g. Antanoudiou et al., 2020). However, manipulations that decrease the activity of PV+ neurons typically disrupt gamma oscillations (Fuchs et al., 2007; Sohal., 2012; Antonoudiou et al., 2020), suggesting that a simple disinhibition of principal neurons is unlikely to be a sufficient explanation.

This view is supported by the actions of bicuculline in the current study. These experiments were included to compare the effects of tianeptine’s inhibition of GABA release with the direct effects of GABA_A_ receptor blockade. The preliminary dose-response analysis of bicuculline’s effects on evoked and spontaneous hippocampal activity revealed that doses at and above 0.1 mM resulted in multiple population spikes after stimulation and large-amplitude LFP spikes. This pattern has been observed in previous studies of GABA_A_R blockade (e.g. Suzuki & Smith, 1998; Bragin et al., 2009) and is reminiscent of the interictal spikes observed in epilepsy and epilepsy models (Lai et al., 2023). Epileptiform activity was not observed at a dose of 30 µM bicuculline, but this dose resulted in a pronounced reduction in paired-pulse inhibition, comparable to that observed after tianeptine administration (cf. Leung et al., 2008). However, this dose did not cause an increase in beta or gamma power, instead causing a modest decrease in LFP power across the frequency range. A previous study has reported that local infusion of picrotoxin in the ventral hippocampus of isoflurane-anaesthetised rats causes an increase in LFP power below 20 Hz, with a peak increase at <10 Hz, particularly during the ‘burst’ state, one of the distinct LFP states characteristic of this anaesthetic (Gwilt et al., 2020). Although there are several differences in methodology, this pattern is reminiscent of the spectral changes observed after higher doses of bicuculline in the current study, but does not mirror the increase in rhythmic beta and gamma activity observed after tianeptine administration. This suggests that tianeptine’s LFP signature cannot be explained solely by a general disinhibition of principal neurons but is dependent on the specific cellular and subcellular targets of the drug. This is perhaps not surprising considering the role of precisely timed interactions between interneurons and pyramidal cells in the generation of gamma rhythms.

As in our previous study of tianeptine’s effects on LFP activity (Burt et al., 2025), the in vivo components of the current study were carried out under urethane anaesthesia. This removes the complicating effects of tianeptine’s locomotor-stimulant actions (see Han et al., 2022; Trigo et al., 2022) at the expense of a modified baseline of excitatory and inhibitory transmission. Urethane potentiates the activation of GABA_A_ receptors but inhibits AMPA and NMDA receptors (Hara & Harris, 2002). Consequently, long-term potentiation (LTP) of synaptic strength requires more intense tetanisation under urethane anaesthesia, but can still be readily induced (Riedel et al., 1994). Anaesthetics that potentiate GABA_A_R activation can enhance PPI of the CA1 population spike in vivo (Pearce et al., 1989). Surprisingly, however, urethane reduces PPI, perhaps owing to its actions on excitatory neurotransmission (Shirasaka & Wasterlain 1995). Regarding LFP activity in the presence of urethane, the pattern is similar to that observed during sleep, characterised by a slow-wave-sleep-like state dominated by large-amplitude irregular activity (LIA), and periodic transitions into a theta-like state reminiscent of REM sleep (Pagliardini et al., 2013; Martin et al., 2019). Beta and gamma oscillations are readily induced under urethane anaesthesia (e.g. Penttonen et al., 2001; Martin, 2001), and our preliminary results indicate that tianeptine causes beta / low-gamma oscillations in freely moving mice (Trigo et al., 2022), ruling out the possibility that this activity is an artifact of anaesthesia in the case of tianeptine.

Both male and female rats were used for the in vivo experiments reported here, and although the numbers are insufficient for a statistical analysis of sex differences, tianeptine’s actions on PPI and LFP activity were qualitatively and quantitatively similar in both female and male rats. This is consistent with our preliminary observations of similar beta-frequency oscillations in freely moving female and male mice following systemic tianeptine administration (Trigo et al., 2022). However, it is possible or even likely that sex differences in tianeptine’s actions may be found in some brain regions if sufficiently powered studies are conducted. For example, sex differences in mouse µ-OR expression have been reported in several brain areas including the periaqueductal grey (Loyd et al., 2008), anterior cingulate cortex and somatosensory areas (Zamfir et al., 2023), and the amygdala (Won et al., 2023); in the latter study, a substantially higher proportion of PV-positive neurons was observed in the amygdala of female compared to male mice; however no difference was observed in the hippocampus.

In summary, our key result, obtained using both in vitro and in vivo techniques, is that tianeptine causes a reduction in hippocampal GABAergic transmission, disinhibiting CA1 pyramidal cells. This ability is shared by other µ-OR agonists. Together with previous evidence that tianeptine can potentiate AMPA-receptor-mediated synaptic transmission via an interaction with postsynaptic signalling pathways (Zhang et al., 2013; Mariano et al., 2026), this raises the possibility that tianeptine enhances excitatory neurotransmission via simultaneous and complementary effects on both sides of the synapse—limiting presynaptic GABA release while enhancing postsynaptic depolarisation.

## Acknowledgements

This work was supported by an Alzheimer’s Research UK (ARUK) Scotland Network Centre small grant awarded to Stephen Martin and Szu-Han Wang. We would like to thank the staff of the Medical Sciences Resource Unit for animal husbandry and welfare, and Patrick Spooner for LabView programming.

